# SARS-CoV-2 infection in the lungs of human ACE2 transgenic mice causes severe inflammation, immune cell infiltration, and compromised respiratory function

**DOI:** 10.1101/2020.07.09.196188

**Authors:** Emma S. Winkler, Adam L. Bailey, Natasha M. Kafai, Sharmila Nair, Broc T. McCune, Jinsheng Yu, Julie M. Fox, Rita E. Chen, James T. Earnest, Shamus P. Keeler, Jon H. Ritter, Liang-I Kang, Sarah Dort, Annette Robichaud, Richard Head, Michael J. Holtzman, Michael S. Diamond

**Author notes:** Corresponding author: Michael S. Diamond, M.D., Ph.D. Contributed equally (Emma S. Winkler, Adam L. Bailey).

## Abstract

Severe Acute Respiratory Syndrome Coronavirus -2 (SARS-CoV-2) emerged in late 2019 and has spread worldwide resulting in the Coronavirus Disease 2019 (COVID-19) pandemic. Although animal models have been evaluated for SARS-CoV-2 infection, none have recapitulated the severe lung disease phenotypes seen in hospitalized human cases. Here, we evaluate heterozygous transgenic mice expressing the human ACE2 receptor driven by the epithelial cell cytokeratin-18 gene promoter (K18-hACE2) as a model of SARS-CoV-2 infection. Intranasal inoculation of SARS-CoV-2 in K18-hACE2 mice results in high levels of viral infection in lung tissues with additional spread to other organs. Remarkably, a decline in pulmonary function, as measured by static and dynamic tests of respiratory capacity, occurs 4 days after peak viral titer and correlates with an inflammatory response marked by infiltration into the lung of monocytes, neutrophils, and activated T cells resulting in pneumonia. Cytokine profiling and RNA sequencing analysis of SARS-CoV-2-infected lung tissues show a massively upregulated innate immune response with prominent signatures of NF-kB-dependent, type I and II interferon signaling, and leukocyte activation pathways. Thus, the K18-hACE2 model of SARS-CoV-2 infection recapitulates many features of severe COVID-19 infection in humans and can be used to define the mechanistic basis of lung disease and test immune and antiviral-based countermeasures.

## INTRODUCTION

Severe Acute Respiratory Syndrome Coronavirus-2 (SARS-CoV-2) is the recently emerged RNA virus responsible for the Coronavirus Disease 2019 (COVID-19) pandemic. Clinical disease is variable, ranging from asymptomatic infection to multi-organ failure and death, with a case-fatality rate of ∼5%. The binding of the SARS-CoV-2 spike protein to human angiotensin-I converting enzyme-2 (hACE2) targets the virus to type II pneumocytes within the lung, resulting in injury, inflammation, and subsequent respiratory distress^1,2^. Other COVID-19 manifestations (*e.g*. cardiac dysfunction, coagulopathy, and gastrointestinal tract symptoms) suggest that extra-pulmonary sites of infection contribute to disease pathogenesis in some patients^3^.

The development of countermeasures that reduce COVID-19 morbidity and mortality is a priority for the global research community, and animal models are essential for this effort. Although several animal species used in laboratory research have been evaluated for susceptibility to SARS-CoV-2 infection, none have recapitulated the severe disease seen in hospitalized human cases. Hamsters, ferrets, and non-human primates develop mild to moderate viral disease and recover spontaneously^4,5^. Conventional laboratory strains of mice cannot be infected efficiently by SARS-CoV-2 because hACE2 but not mouse ACE2 supports SARS-CoV-2 binding^6,7^. Multiple strategies for introducing hACE2 into mice have been developed including (1) transient introduction of hACE2 via adenoviral viral vectors^8^, (2) expression of hACE2 as a transgene driven by heterologous gene promoters^9,10^, or (3) expression of hACE2 by the mouse ACE2 promoter^11,12^. While these animals all support SARS-CoV-2 infection, none cause severe disease or lethality. Thus, an animal model is still urgently needed for understanding the biology of severe SARS-CoV-2 infection and evaluating the efficacy of countermeasures for COVID-19.

The K18-hACE2 transgenic (K18-hACE2) mice, in which hACE2 expression is driven by the epithelial cell cytokeratin-18 (K18) promoter^13^, were originally developed to study SARS-CoV pathogenesis and cause lethal infection^9^. Here, we evaluate heterozygous hACE2 transgenic mice as a model for severe COVID-19 disease. After intranasal SARS-CoV-2 inoculation, K18-hACE2 mice rapidly lost weight starting at 4 days post infection (dpi) and began to succumb to disease at 7 dpi. High levels of viral RNA and infectious virus were detected in the lungs of infected animals at 2, 4, and 7 dpi by RT-qPCR, *in situ* hybridization, and plaque forming assays. Infection was accompanied by declines in multiple parameters of pulmonary function, substantial cellular infiltrates in the lung composed of monocytes, neutrophils, and activated T cells, high levels of pro-inflammatory cytokines and chemokines in lung homogenates, and severe interstitial and consolidative pneumonia. Because of its severe disease and intense immune cell infiltration, the K18-hACE2 model of SARS-CoV-2 infection may facilitate evaluation of immunomodulatory and antiviral drugs against COVID-19 and our understanding of immune-mediated mechanisms of pathogenesis.

## RESULTS

### K18-hACE2 mice are highly susceptible to SARS-CoV-2 infection

We inoculated 8-week-old heterozygous K18-hACE2 mice of both sexes via intranasal route with 2.5×10^4^ PFU of SARS-CoV-2 (strain 2019n-CoV/USA_WA1/2020). Beginning at 4 days post-infection (dpi), K18-hACE2 mice demonstrated marked weight loss, and by 7 dpi most animals had lost approximately 25% of their body weight (**Fig 1a**), with many becoming moribund. High levels of infectious SARS-CoV-2 (**Fig 1b**) and viral RNA (**Fig 1c**) were detected in lung homogenates at 2, 4, and 7 dpi, whereas lower levels were present in other tissues (*e.g*., heart, spleen, kidney). Virtually no viral RNA was measured in gastrointestinal tract tissues or in circulation until 7 dpi in the serum and colon, and this was only in a subset of animals (**Fig 1d**). The tissues supporting SARS-CoV-2 infection in this model mirrored the pattern of hACE2 expression, with the highest receptor levels in the lungs, colon, kidney, and brain (**Extended Data Fig 1a**). Levels of hACE2 declined in the lung over the course of infection (**Extended Data Fig 1b**), suggesting either receptor downregulation, hACE2 shedding, or death of hACE2-expressing cells after SARS-CoV-2 infection. A subset of infected K18-hACE2 mice had high levels of viral RNA and infectious virus in the brain (**Fig 1d-e**), consistent with previous reports with SARS-CoV and SARS-CoV-2 in hACE2 transgenic mice^9,12,14^. As no infectious virus and only low levels of viral RNA were detected in the brain of the majority (60%) of animals, the observed clinical disease is more consistent with lung and not brain infection. Staining for viral RNA in brain tissue by *in situ* hybridization showed that only one of six animals was positive at 7 dpi.; this animal had disseminated infection throughout the cerebral cortex with noticeable sparing of the olfactory bulb and cerebellum (**Extended Data Fig 2**).

**Figure 1.**
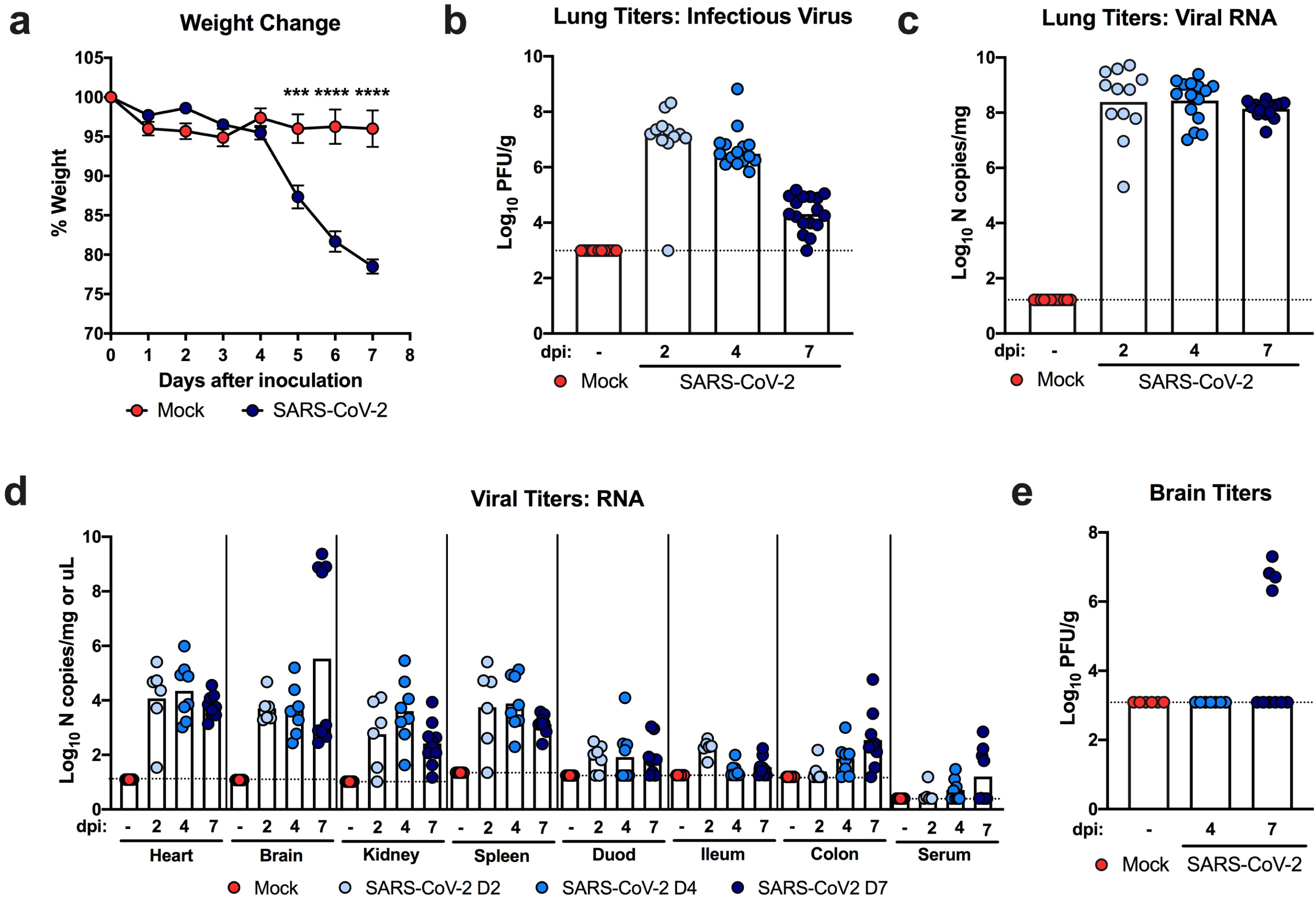
SARS-CoV-2 infection in K18-hACE2 mice. Eight to nine-week-old male and female K18-hACE2 transgenic mice were inoculated via the intranasal route with 2.5 × 10^4^ PFU of SARS-CoV-2. **a**. Weight change was monitored (two experiments, n = 10; two-way ANOVA: *** *P* < 0.001, **** *P* < 0.0001, symbols represent the mean ± SEM). **b-c**. Viral burden in the lungs was analyzed at 2, 4 and 7 dpi by plaque assay for infectious virus (**b**) and qRT-PCR for viral RNA levels (**c**). **d**. Viral RNA levels in indicated tissues (heart, brain, kidney, spleen, serum, and gastrointestinal tract) at 2, 4, and 7 dpi as measured by qRT-PCR. **e**. Viral burden in the brains as measured by plaque assay (two experiments, n = 10). For **b-e**, bars represent the mean and the dotted line indicates the limit of detection.

### Histopathological changes in the lung after SARS-CoV-2 infection

Analysis of hematoxylin and eosin-stained lung sections from K18-hACE2 mice infected with SARS-CoV-2 (**Fig 2a**) showed a progressive inflammatory process. At 2 dpi, we observed accumulation of immune cells confined predominantly to perivascular sites. By 4 dpi, these immune cell infiltrates involved a greater area of the lung with focal collections into adjacent alveolar spaces with alveolar wall thickening. By 7 dpi, immune cells, including neutrophils and mononuclear cells were found throughout the lung in alveolar and interstitial locations along with interstitial edema and consolidation. To correlate histopathological findings with sites of SARS-CoV-2 infection, we also stained lung sections for viral RNA using *in situ* hybridization (**Fig 2b**). At 2 dpi, expression of SARS-CoV-2 RNA was localized predominantly to alveolar epithelial cells and a few airway epithelial cells. This pattern also was seen at 4 dpi, but with more diffuse spread throughout the lung. By 7 dpi, the level of viral RNA expression was diminished and associated with cellular debris and collapsed alveoli. No significant viral RNA signal was localized to immune cells. Together, these findings provide evidence of a progressive and widespread viral pneumonia with perivascular and pan-alveolar inflammation characterized by immune cell infiltration, edema, and lung consolidation.

**Figure 2.**
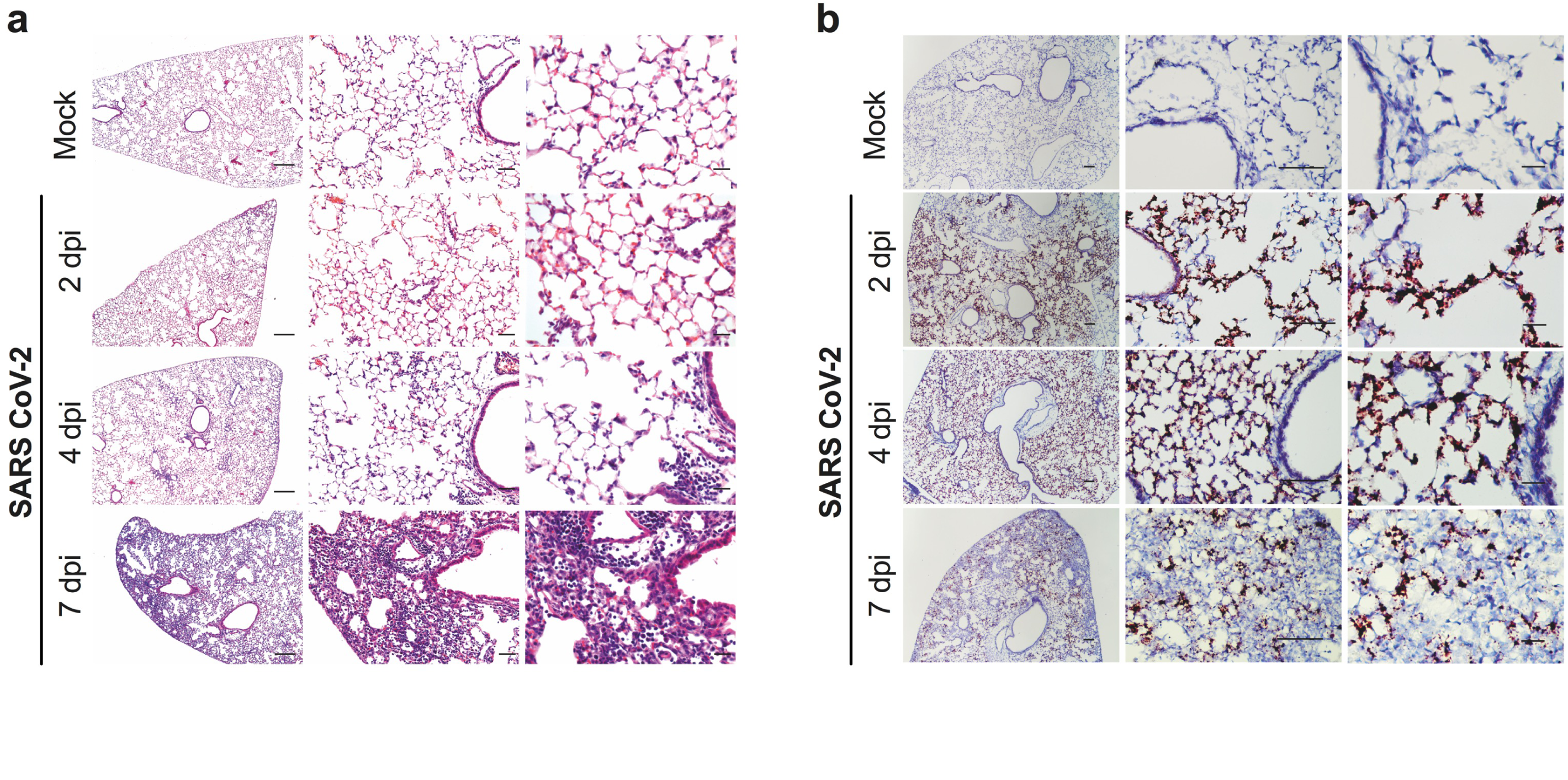
Histopathological analysis of SARS-CoV-2 infection in K18-hACE2 mice. **a**. Hematoxylin and eosin staining of lung sections from K18-hACE2 mice following mock infection or after intranasal infection with 2.5 × 10^4^ PFU of SARS-CoV-2 at 2, 4, and 7 dpi. Images show low- (left; scale bars, 250 μm), medium- (middle; scale bars, 50 μm), and high-power magnification (right; scale bars, 25 μm). Representative images from n = 6 per group. **b**. SARS-CoV-2 RNA *in situ* hybridization of lung sections from K18-hACE2 mice for conditions in (**a**). Images show low- (left; scale bars, 100 μm), medium- (middle; scale bars, 100 μm), and high-power magnification (right; scale bars, 10 μm). Representative images from n = 6 per group.

### Extra-pulmonary histopathology after SARS-CoV-2 infection

We examined additional tissues implicated in the pathogenesis of severe COVID-19 in humans, including the brain, heart, liver, kidney, and spleen. Brain tissues of K18-hACE2 mice with minimal detectable SARS-CoV-2 infection appeared normal, whereas the one brain with a high level of infection at 7 dpi showed multiple foci of inflammatory cells (*e.g*., neutrophils, lymphocytes, and monocytes) involving the meninges, the subarachnoid space, parenchymal blood vessels, and the brain parenchyma (**Extended Data Fig 3a**). Abnormalities were observed in 2 of 9 hearts at 4 dpi (*e.g*., scattered hypereosinophilic cardiomyocytes with pyknotic nuclei) and most livers at 4 and 7 dpi (*e.g*., areas of inflammatory cell infiltrates and hepatocyte loss) (**Extended Data Fig 3b-c**). In one kidney at 4 dpi, we observed focal acute tubular injury (**Extended Data Fig 3d**); otherwise, the kidneys showed no apparent abnormalities. The spleen in SARS-CoV-2-infected K18-hACE2 mice appeared normal (**Extended Data Fig 3e**), and fibrin thrombi were not detected in any of the extra-pulmonary organs examined.

### Pathophysiology of SARS-CoV-2 infection

To assess for clinically-relevant changes in physiology over the course of SARS-CoV-2 infection in K18-hACE2 mice, we measured clinical chemistry and hematological parameters from peripheral blood samples (**Fig 3**). Plasma levels of sodium, potassium, and chloride concentrations and the anion gap all trended slightly downward at 7 dpi (**Fig 3a-d)** whereas plasma bicarbonate noticeably increased (**Fig 3e**), possibly as a result of poor gas exchange resulting from lung pathology or decreased respiratory drive. Other plasma analytes including calcium, glucose, and blood urea nitrogen were unchanged (**Fig 3f-h**). Hematocrit and plasma hemoglobin levels increased later in the course of infection, possibly because of reduced water intake and hemoconcentration (**Fig 3i-j**). We also observed a modest prolongation in the prothrombin time at 7 dpi that was preceded by an increase in D-dimer concentrations on 2 and 4 dpi (**Fig 3k-l**).

**Figure 3.**
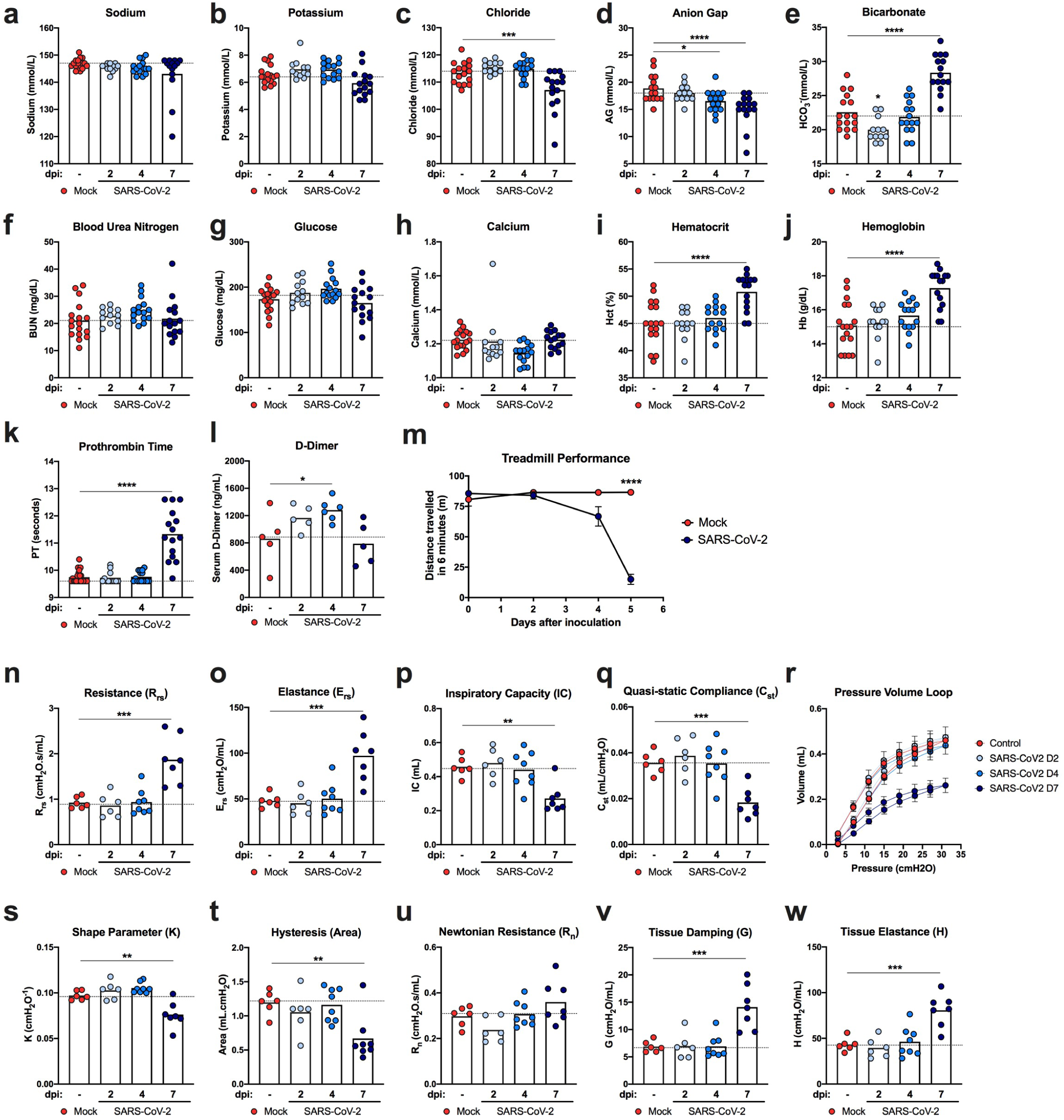
Functional consequences of SARS CoV-2 infection in K18-hACE2 mice. **a-k**. Clinical chemistry and hematological parameters from the peripheral blood of mock-treated or SARS-CoV-2 infected animals at 2, 4, and 7 dpi (two experiments, n = 13-16 per group; one-way ANOVA with Dunnett’s test; * *P* < 0.05; *** *P* < 0.001; **** *P* < 0.0001, bars represent the mean, and the dotted line indicates the mean value of the mock-treated group). **l**. Serum D-dimer levels of mock-treated or SARS-CoV-2 infected animals at 2, 4, and 7 dpi (two experiments, n = 5 per group; one-way ANOVA with Dunnett’s test; * *P* < 0.05; *** *P* < 0.001; **** *P* < 0.0001, bars represent the mean, and the dotted line indicates the mean value of the mock-treated group). Asterisks indicate statistical significance compared to mock infection. **m**. Results of a treadmill performance test as measured by the distance traveled in 6 minutes. (two experiments, n = 10; two-way ANOVA: **** *P* < 0.0001, symbols represent the mean ± SEM). **n-w**. Respiratory mechanics parameters from the lung function assessment in mock-treated or SARS-CoV-2 infected male and female mice at 2, 4, and 7 dpi. Individual results with group mean are shown. **n**. Inspiratory capacity. **o**. Respiratory system resistance. **p**. Respiratory system elastance. **q**. Pressure-Volume (PV) loops. **r**. Static compliance. **s**. Shape parameter K. **t**. Hysteresis (Area). **u**. Newtonian resistance. **v**. Tissue damping. **w**. Tissue elastance (two experiments, n = 6-7 per group; one-way ANOVA with Dunnett’s test; * *P* < 0.05; *** *P* < 0.001; **** *P* < 0.0001, bars represent the mean and the dotted line indicates the mean value of the mock-treated group).

We next examined the impact of SARS-CoV-2 infection on pulmonary and cardiac function using a treadmill stress-test to assess exercise tolerance (**Fig 3m**). Compared to mock-infected controls, at 4 dpi, a subset of SARS-CoV-2-infected K18-hACE2 mice began to show reduced exercise tolerance, as measured by decreased distance travelled. However, by 5 dpi, all infected K18-hACE2 mice had substantially reduced exercise tolerance compared to mock-infected animals or their own pre-infection baseline performance (**Fig 3m**).

To examine changes to the biophysical properties of the lung over the course of SARS-CoV-2 infection, we mechanically ventilated mice via tracheostomy and performed several forced-oscillation tests to determine various respiratory mechanics parameters (**Fig 3n-w**). Infected animals showed normal lung biomechanics at 2 and 4 dpi but had markedly abnormal values in most parameters at 7 dpi relative to mock-infected controls. These abnormalities included reduced inspiratory capacity as well as increased respiratory system resistance and elastance (**Fig 3n-p**). Collectively, these changes resulted in a downward deflection of the pressure-volume loop (**Fig 3r**) with concomitant decreases in static compliance (**Fig 3q**), the shape-describing K parameter (**Fig 3s**), and loop hysteresis (**Fig 3t**), which together indicate reduced lung compliance and distensibility. Further analysis using broadband forced oscillation maneuvers^15^ revealed that SARS-CoV-2 infected mice at 7 dpi had relatively normal Newtonian resistance (**Fig 3u)**, which is primarily a reflection of resistance in larger conducting airways. In contrast, mice at 7 dpi had marked increases in tissue damping (**Fig 3v**) and elastance (**Fig 3w**); these parameters measure the dissipation and storage of oscillatory energy in parenchymal tissue and reflect tissue and peripheral airway resistance and elastic recoil (*i.e*., tissue stiffness), respectively. The measurements of mechanical properties of the respiratory system suggest that SARS-CoV-2 infection in K18-hACE2 mice predominantly causes disease in the alveoli and lung parenchyma, and not in the conducting airways, which is consistent with both our histopathological analysis in mice and measurements of pulmonary function in humans with viral pneumonia and respiratory failure including COVID-19^16^.

### The immune response to SARS-CoV-2 Infection in the lungs

An excessive pro-inflammatory host response to SARS-CoV-2 infection is hypothesized to contribute to pulmonary pathology and the development of respiratory distress in some COVID-19 patients^17^. To evaluate the composition of the immune cell response in SARS-CoV-2-infected K18-hACE2 mice, we performed flow cytometric analysis on lung homogenates and bronchoalveolar lavage (BAL) fluid at three time points after intranasal virus inoculation (**Fig 4a-b, Extended Data Fig 4**). Consistent with our histopathological analysis, we observed increased numbers of CD45^+^ immune cells in the BAL beginning at 2 dpi and in the lung at 4 dpi. The cellular infiltrates at 4 and 7 dpi in the lung were composed principally of myeloid cell subsets including Ly6G^+^ neutrophils, Ly6C^+^ monocytes, and CD11b^+^CD11c^+^ dendritic cells. In the BAL fluid, monocyte numbers peaked at 4 dpi, and levels of neutrophils and dendritic cells continued to rise through 7 dpi. Accumulation of monocytes in the BAL fluid coincided with a decrease in the number of tissue-resident alveolar macrophages, an observation consistent with scRNA-seq analysis of BAL fluid of patients with severe COVID-19 disease^18,19^. By 7 dpi, we also observed an increase in several lymphoid cell subsets in the lung including NK1.1^+^ natural killer cells, γδ CD3^+^ T cells, CD3^+^CD4^+^ T cells, CD3^+^CD8^+^ T cells, and activated CD44^+^CD3^+^CD8^+^ T cells (**Fig 4a**).

**Figure 4.**
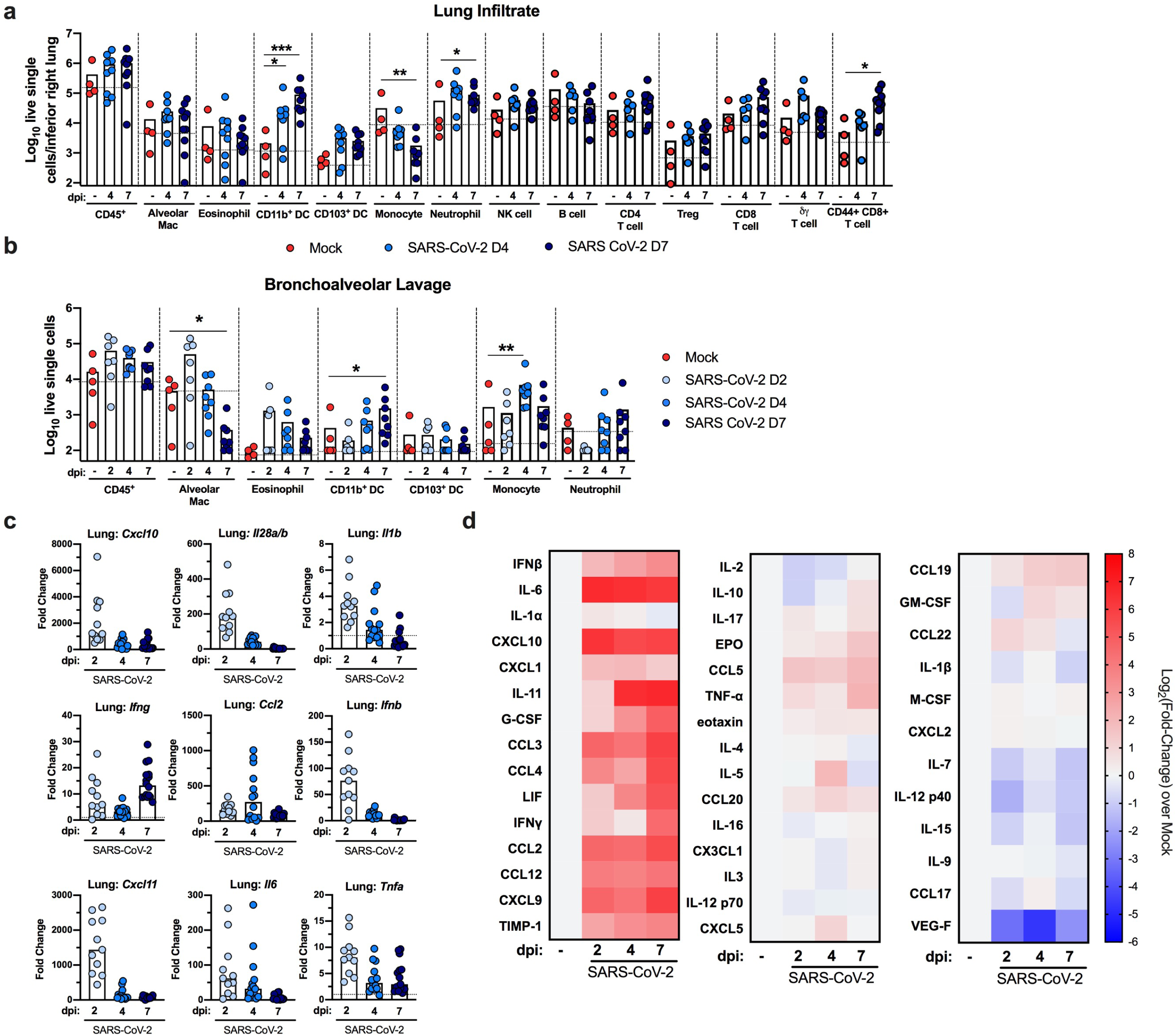
The immune response to SARS-CoV-2 Infection in the lungs of K18-hACE2 mice. **a-b**. Flow cytometric analysis of lung tissues (**a**) and bronchoalveolar lavage (**b**) at 2, 4, and 7 dpi post-SARS-CoV-2 infection (two experiments, n = 4-6 per group; one-way ANOVA; * *P* < 0.05; ** *P* < 0.01; *** *P* < 0.001, bars represent the mean and the dotted line indicates the mean value of the mock-treated group). Asterisks indicate statistical significance compared to mock infection. **c**. Fold change in gene expression of indicated cytokines and chemokines as determined by RT-qPCR, normalized to *Gapdh*, and compared to naïve controls in lung homogenates at 2, 4 and 7 dpi (two experiments, n = 9-11 per group). Dotted line indicates the average level of cytokine or chemokine transcript in naïve mice. **d**. Heat-maps of cytokine levels as measured by multiplex platform in lung tissue of SARS-CoV-2-infected mice at 2, 4, and 7 dpi. For each cytokine, fold-change was calculated compared to mock-infected animals and Log2(fold-change) was plotted in the corresponding heat-map (two experiments, n = 9-11 per group, associated statistics are reported in **Extended Data Fig 5**).

Extensive changes in cytokine profiles are associated with COVID-19 disease progression^20-22^. Compared to the lungs of uninfected K18-hACE2 control mice, we observed induction of *Ifnb, Il28, Ifng, Cxcl10, Cxcl11*, and *Ccl2* mRNA over the first week (**Fig 4c**) with highest expression occurring at 2 dpi for all cytokines except *Ifng* and *Ccl2*. We also measured protein levels in the lungs using a multiplex assay of 44 different cytokines and chemokines (**Fig 4d, Extended Data Fig 5**). Although mRNA expression was highest at 2 dpi, almost all up-regulated pro-inflammatory cytokines (IFNβ, IL-6, CXCL10, CXCL9, CCL5, CCL12, TIMP-1, TNFα, and G-CSF), T cell-associated cytokines (IL-10, IFNγ, and IL-2), and myeloid cell-associated chemokines (CCL2, CCL3, CCL4, CXCL1, and LIF) peaked at 7 dpi. These data are consistent with cytokine profiling of serum from human COVID-19 patients and transcriptional analysis of the BAL fluid of human patients, which showed that elevated levels of IL-10, IL-6, IL-2, IL-7, G-CSF, CXCL10, CCL2, CCL3, and TNF-α correlate with disease severity^19,23-25^. Overall, our data suggest that in the context of the inflammatory response to SARS-CoV-2 in the lungs of K18-hACE2 mice, many cytokines and chemokines are induced, with some having sustained expression and others showing rapid up-and down regulation patterns.

### Distinct transcriptional signatures are associated with early and late immune responses to SARS-CoV-2 infection

Studies in other small animals and humans have reported cytokine signatures coupled with delayed type I interferon (IFN) signaling or elevated IFN signatures in the lung^26,27^. To assess how the kinetics of infection and ensuing inflammation modulate the cytokine and IFN response to SARS-CoV-2, we performed RNA sequencing of lung homogenates of K18-Tg mice at 0 (mock), 2, 4, and 7 dpi. Principal component analysis (PCA) revealed distinct transcriptional signatures associated at 7 dpi (**Fig 5a**) with overlapping signatures at 2 and 4 dpi. Hundreds of genes were differentially expressed at all time points compared to mock-infected animals (**Fig 5b**), many of these associated with IFN signaling, NF-kB-dependent cytokine responses, or leukocyte activation. In agreement with the PCA, only 449 differentially expressed genes were shared at all time points when compared to mock. In contrast, 1,975 unique differentially expressed genes were identified between mock to 7 dpi whereas only 59 and 152 genes were different between mock and 2 and 4 dpi, respectively (**Fig 5c**). Gene ontology analysis of the top upregulated genes at all time points showed enrichment of gene clusters in cytokine-mediated signaling, type I and II IFN signaling, neutrophil activation, and pathogen recognition receptor signaling (**Fig 5d**). Upregulation of gene sets involved in cytokine-mediated signaling, neutrophil activation, cellular responses to type II IFN, and toll-like receptor signaling were most pronounced at 7 dpi (**Fig 5e-g, Extended Data Fig 6a-c, Supplementary Table 1**). Of note, different genes in the type I IFN signaling pathway were upregulated at 2 and 4 dpi (*e.g*., *Irf9, Irf7, Stat1*, and certain IFN-stimulated genes (ISGs) *Isg15, Mx1, Oas3, Ifit1, Ifit2, and Ifit3*) versus 7 dpi (*e.g*., *Ifnar1/2, Tyk2, Irf1* and certain ISGs *Samhd1, Oas2, and Ifitm1*). This suggests a temporally distinct type I IFN response (**Fig 5f**), which has been described previously with IFN*α* and IFN*β* subtypes^28-30^. Alternatively, the differences in IFN and ISG signatures at early and late time points could reflect differential signaling contributions of type I and III IFNs, as these cytokines are both expressed in the lung after SARS-CoV-2 infection^31^ and induce overlapping yet non-identical sets of ISGs^32^. Collectively, the RNA sequencing data from the lungs of K18-hACE2 mice show distinct immune signatures associated with early infection (days 2 and 4) and late (day 7) SARS-CoV-2 infection.

**Figure 5.**
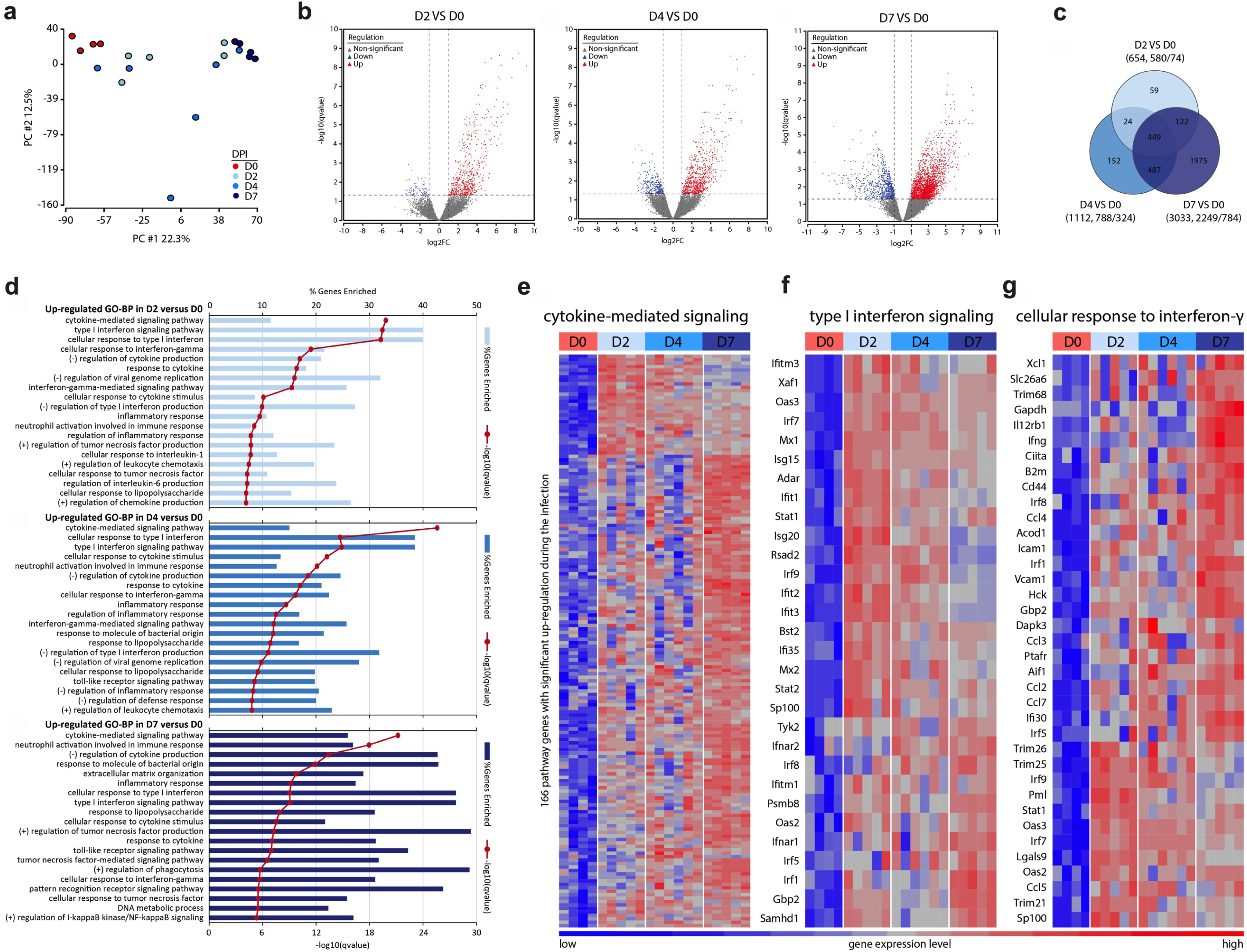
Distinct transcriptional signatures are associated with early and late immune responses to SARS-CoV-2 infection. RNA sequencing analysis from the lung homogenates of naive K18-hACE2 mice and at 2, 4, and 7 dpi (two experiments, n = 4-6 group) **a**. Principal component analysis performed for 20 samples with the log2-transformed gene-level counts per million (log2cpm) data **b**. Volcano plots comparing differentially-expressed genes from samples taken at day 2 versus day 0, day 4 versus day 0, and day 7 versus day 0. Red and blue indicate upregulated (red) and downregulated (blue) genes that demonstrated a fold-change > 2 and false discovery rate (q-value) < 0.05. The dashed horizontal lines mark a q-value of 0.05 and vertical lines indicate log2 fold-change of 1. Each dot in the volcano plots represents a single gene. **c**. Venn diagram of overlapping genes identified in differential expression analysis when comparing mock to 2, 4, and 7 dpi. Numbers in the parenthesis under each comparison indicates the total number of significantly differential genes followed by the proportion of the total that are up and down-regulated. **d**. GO Enrichment Analysis of biological process terms enriched in up-regulated genes from comparisons of mice 2, 4, and 7 days dpi against mock-infected mice. Terms were ranked by the false discovery rate (q-value), and the top 20 are listed after eliminating redundant terms. **e-g**. Heat maps of significantly up-regulated genes during SARS-CoV-2 infection enriched in cytokine-mediated signaling pathway (**e**), type I IFN signaling pathway (**f**), and cellular response to IFNγ (**g**) identified through Gene Ontology analysis. Genes shown in each pathway are the union of the differentially expressed genes from the three comparisons (2, 4, and 7 dpi versus mock-infected). Columns represent samples and rows represent genes. Gene expression levels in the heat maps are z score-normalized values determined from log2cpm values.

## DISCUSSION

In this study, we found that SARS-CoV-2 infection of K18-hACE2 transgenic mice causes severe pulmonary disease. After intranasal SARS-CoV-2 inoculation, K18-hACE2 mice rapidly lost weight after 4 dpi and began to succumb to disease at 7 dpi. High levels of viral RNA and infectious virus were detected in the lungs of infected animals at 2, 4, and 7 dpi by RT-qPCR, *in situ* hybridization, and plaque assay. Infection was accompanied by high levels of pro-inflammatory cytokines and chemokines in the lung and an impressive cellular infiltrate comprised primarily of monocytes, neutrophils, and T cells. The combined infection and inflammation resulted in severe interstitial pneumonia characterized by collapsed alveolar spaces. This caused detrimental changes in lung physiology including decreased exercise tolerance, reduced inspiratory capacity, and stiffening of the lung parenchyma.

SARS-CoV-2 infection is subclinical or mild in most human cases. A small, yet clinically important fraction develop life-threatening disease requiring hospitalization and intensive care. Mild disease is a feature of SARS-CoV-2 infection in naturally susceptible animals including hamsters, ferrets, cats, and non-human primates^5^. This is perhaps unsurprising given that the strongest risk factors for developing severe COVID-19 in humans (*e.g*., old age, cardiovascular disease, and diabetes) are absent in many laboratory animals. Mild to moderate disease is seen in many rodent models of SARS-CoV-2 infection, including those expressing hACE2 via viral vectors or transgenes^8,11,12,33^. Thus, the severity of disease we observed following SARS-CoV-2 infection of K18-hACE2 mouse is unique. As the onset of severe clinical disease in K18-hACE2 mice occurs days after peak viral infection and is associated with high levels of infiltrating immune cells and inflammatory mediators in the lung, immune responses likely contribute to pathogenesis.

The histopathological changes we observed in the infected lungs of K18-hACE2 mice correlate with the impaired pulmonary function. Pneumocytes become infected early, which led to recruitment of leukocytes into the pulmonary interstitium, production of proinflammatory cytokines, injury to parenchymal cells, collapse of the alveolar space, and compromise of gas exchange, all of which could cause the hypercapnia we observed at 7 dpi. This course is remarkably consistent with human disease in which rapid early viral replication is followed by inflammatory responses, which are believed to contribute to pathology, morbidity, and mortality^34^.

A fundamental understanding of the immunological processes that influence COVID-19 disease is needed to select immunomodulatory interventions that target key cell types or pathways. We saw substantial immune cell accumulation in the lungs of K18-hACE2 mice, an observation consistent with post-mortem analysis of human patients^35^. Infiltrates were composed primarily of myeloid cells including monocytes and neutrophils as well as activated CD8^+^ T cells and corresponded with high levels of chemokines that drive their migration. The lymphopenia associated with severe COVID-19 in humans is attributed in part to the immune cell migration into inflamed tissues^21,36^. In transcriptional analyses of BAL fluid from infected humans with severe COVID-19, an accumulation of CD8^+^ T cells, neutrophils, and monocytes coincided with the loss of alveolar macrophages^18,19^. In our study, using cytokine analysis and RNA-sequencing of lung homogenates, we detected enhanced expression of several myeloid cell chemoattractants (*e.g*., CCL2, CCL3, CCL4, CXCL1, and CXCL10) and other key inflammatory cytokines (TNF*α*, IL-6, and G-CSF) that correlate with COVID-19 disease severity in humans^26,27^. Given these parallel findings, studies in K18-hACE2 mice evaluating the role of specific immune pathways and cell subsets in disease pathogenesis could inform the selection of immunomodulatory agents for severe COVID-19.

The role of type I IFN in SARS-CoV-2 pathogenesis in this model warrants further investigation, as it has been suggested that a dysregulated type I IFN response contributes to excessive immunopathology. Indeed, in SARS-CoV infection, type I IFN signaling appears pro-inflammatory and not antiviral^37^. Our RNA sequencing analysis revealed differences in the type I IFN gene signatures associated with early and late SARS-CoV-2 infection. Given that ISGs can exert diverse functions apart from their antiviral activities, including inflammatory, metabolic, and transcriptional effects^38^, these two “early” and “late” ISG modules may have different functional consequences following SARS-CoV-2 infection. Furthermore, how these temporally distinct programs are induced and regulated remains uncertain and may be the result of cell-type specificity, kinetics, and sensitivity to different type I or III IFN subtypes.

The development of severe disease following SARS-CoV-2 infection is an important feature of the K18-hACE2 model, although the precise reason for this susceptibility compared to other hACE2 transgenic models remains unknown. Potential explanations include a high number of hACE2 transgene insertions (https://www.jax.org/strain/034860) and/or the strength and cellular specificity of the K18 promoter. Human ACE2 is expressed in multiple tissues in the K18-hACE2 mouse^9^, which could allow for SARS-CoV-2 infection of multiple organs. While we detected viral RNA in several tissues, the lung was the most heavily infected and showed the most consistent and severe histopathological changes; these findings were anticipated given the known tropism of SARS-CoV-2 for the respiratory tract and the intranasal route of infection. Moderate levels of viral RNA also were found in the heart, kidney, and spleen, with peak titers at 2 and 4 dpi, whereas levels in gastrointestinal tract tissues (duodenum, ileum, and colon) were lower. In the gastrointestinal tract of K18-hACE2 mice, hACE2 was expressed most abundantly in the colon, which correlated with infection seen at later time points. Although hACE2 is expressed in the gastrointestinal tract in other hACE2-expressing mice, productive infection was observed only upon intragastric inoculation or at early time points following intranasal infection^11,12^.

We observed dichotomous SARS-CoV-2 infection in the brain, with high virus levels in approximately 40% of mice at 7 dpi, and low levels in the remaining 60% of animals. Infection of the brain also was observed in K18-hACE2 mice infected with SARS-CoV^9,14^ but occurred earlier (at 3 to 4 dpi) and more uniformly. Similar to the experiments with SARS-CoV, we did not detect SARS-CoV-2 in the olfactory bulb, which suggests that both SARS-CoV and SARS-CoV-2 cross the blood-brain barrier instead of traversing the cribriform plate and infecting neuronal processes near the site of intranasal inoculation^39^. Notwithstanding this data, and unlike SARS-CoV, alterations in smell and taste are features of SARS-CoV-2 infection in humans^40^, suggesting that cell types within the olfactory system may be susceptible to infection or injury. More study is needed to clarify the routes SARS-CoV-2 dissemination throughout the host and particularly how it accesses the brain in some animals and humans.

While SARS-CoV-2 lung infection in K18-hACE2 mice provides a model for studying severe infection that recapitulates features of COVID-19 in humans, we acknowledge several limitations. The expression of the hACE2 transgene is non-physiological in several respects. It is driven by a non-native (*i.e*., the cytokeratin-18) promotor, resulting in tissue expression levels that are distinct from endogenously-expressed ACE2. ACE2 expression in K18-hACE2 mice is independent of the complex regulatory systems that governs ACE2 levels^41^. As such, comorbid conditions (*e.g*., obesity, hypertension, diabetes) that alter ACE2 expression in humans^41^ likely cannot be modelled faithfully in this transgenic mouse.

In summary, we found that SARS-CoV-2 infection of K18-hACE2 transgenic mice supports robust viral replication in the lung, which leads to severe immune cell infiltration, inflammation, and pulmonary disease. Thus, the K18-hACE2 mouse is an attractive small animal model for defining the mechanisms of the pathogenesis of severe COVID-19 and may be useful for evaluating countermeasures that reduce virus infection or associated pathological inflammatory responses.

## ACKNOWLEDGEMENTS

This study was supported by NIH contracts and grants (75N93019C00062 and R01 AI127828, R01 AI130591, and R35 HL145242) and the Defense Advanced Research Project Agency (HR001117S0019). E. S. W. is supported by T32 AI007163, B.T.M is supported by F32 AI138392, and L.K. is supported by T32 EB021955. We thank Sean Whelan, Susan Cook, and Jennifer Philips for facilitating the studies with SARS-CoV-2 in biosafety level-3, Cathleen Lutz and The Jackson Laboratory for providing mice, Arthur Kim for purifying the CR3022 anti-S mAb, Hana Janova and Matthew Cain for experimental advice, and Robert Schmidt for reviewing a brain histology slides. We also thank David Brunet at SCIREQ Inc. for facilitating use of the flexiVent mouse ventilator.

## AUTHOR CONTRIBUTIONS

A.L.B. and E.S.W. performed the intranasal inoculations of SARS-CoV-2 and clinical analysis. E.S.W., J.M.F., and R.E.C. performed viral burden analysis with support of J.T.E..N.M.K. performed histopathological studies. B.T.M. performed *in situ* hybridization. S.P.K., J.H.R., L.K., and M.J.H. analyzed the tissue sections for histopathology. S.N. and E. S. W. performed immune cell processing for flow cytometry and analysis. A.L.B. performed pulmonary mechanics analysis with training from S.D., and S.D. and A.R. performed analysis of pulmonary mechanics data. A.L.B. and N.M.K. performed treadmill stress-testing analyses. J.Y. and R.H. performed RNA sequencing and analysis. E.S.W. compiled all figures. A.L.B, E.S.W., and M.S.D. wrote the initial draft, with the other authors providing editing comments.

## EXTENDED DATA FIGURE LEGENDS

**Extended Data Figure 1.**
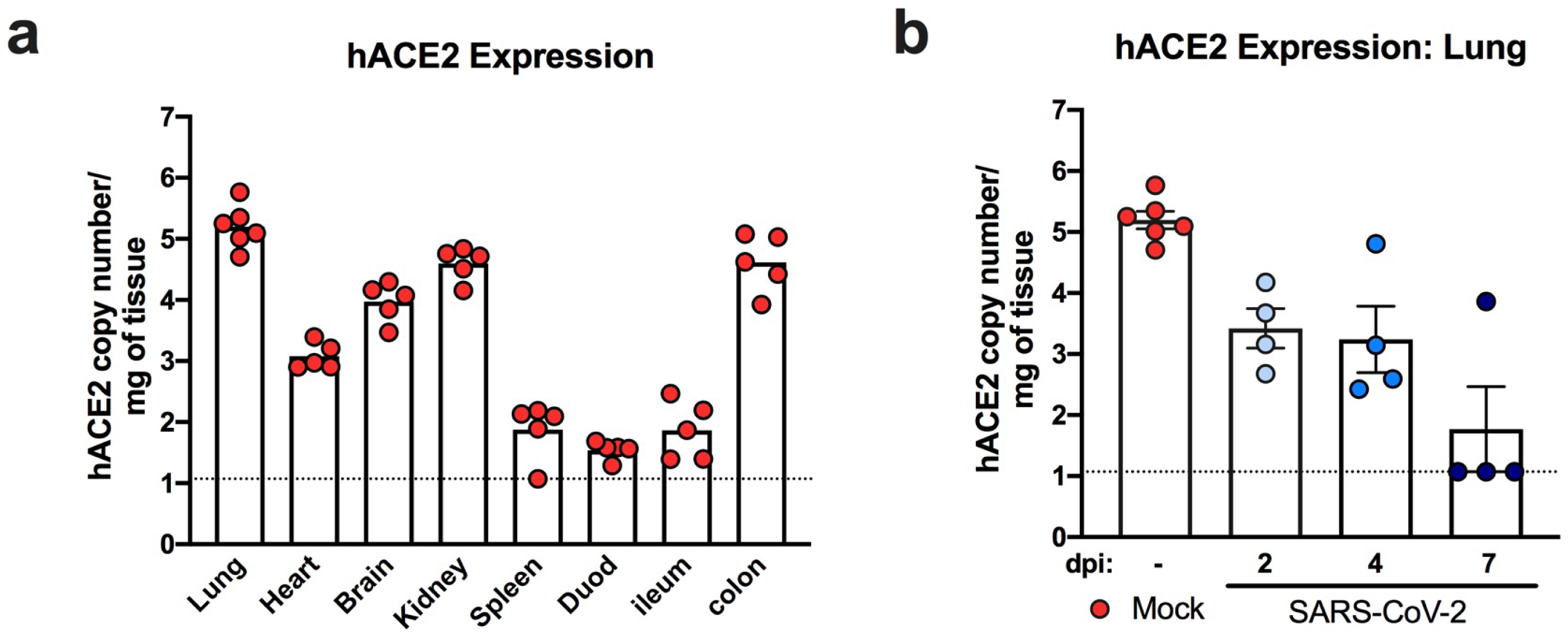
hACE2 expression in the K18-hACE2 model. **a**. mRNA expression levels of hACE2 in the lung, kidney, heart, brain, spleen, duodenum, colon, and ileum of naive K18 hACE2 mice (two experiments, n = 5). **b**. mRNA expression levels of hACE2 in the lungs of K18 hACE2 mice at 2, 4, and 7 dpi following SARS-CoV-2 infection (two experiments, n = 4 per time point, bars represent the mean value of each group).

**Extended Data Figure 2.**
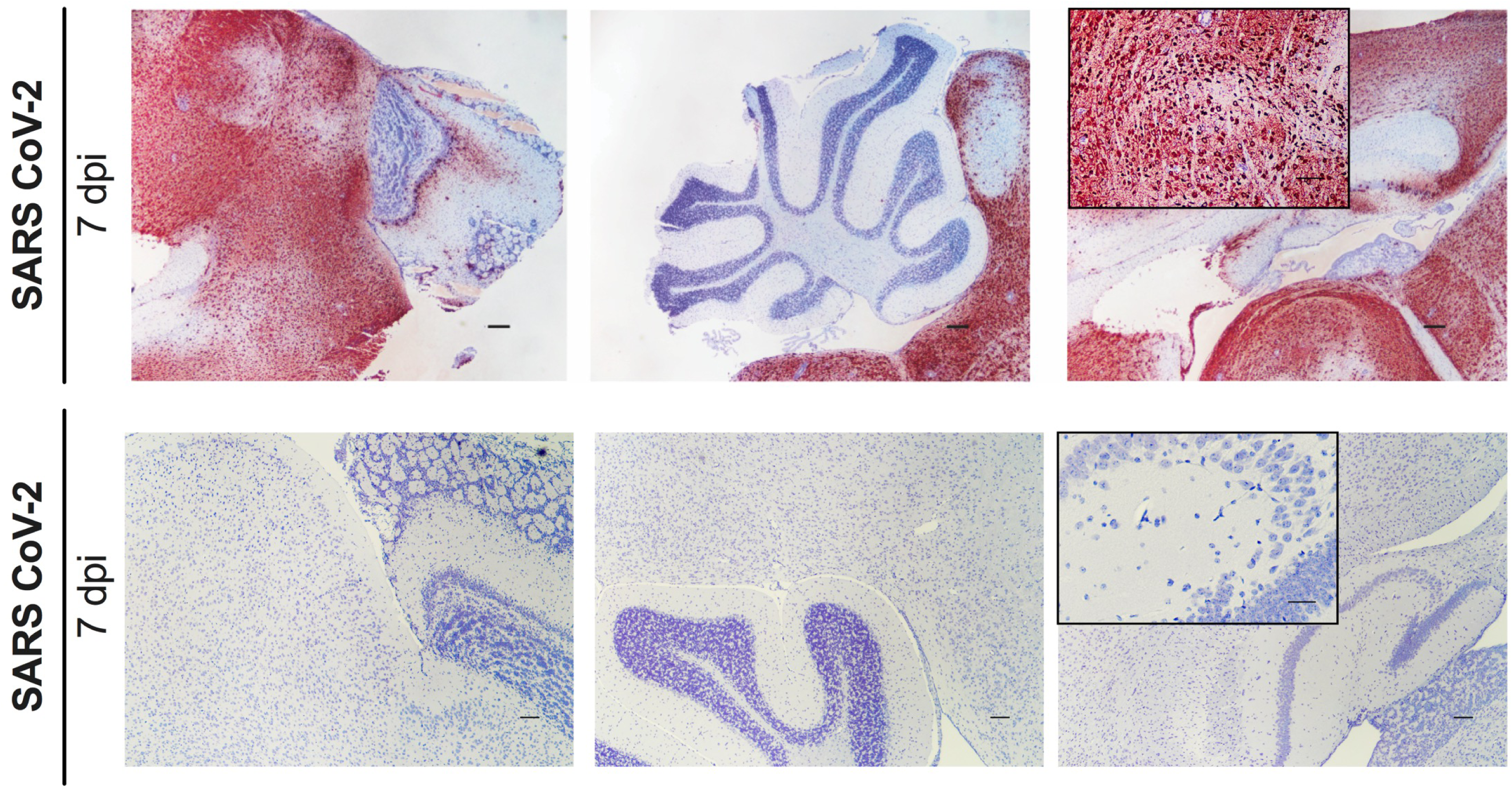
SARS-CoV-2 Infection in the brain. SARS-CoV-2 RNA *in situ* hybridization of brain sections from K18-hACE2 mice following intranasal infection with 2.5 × 10^4^ PFU of SARS-CoV-2 at 7 dpi. Images show low-power magnification (scale bars, 100 μm) with a high-power inset. One of six infected mice stained positively for viral RNA in the brain. Images are from this mouse and another that showed virtually no infection in the brain.

**Extended Data Figure 3.**
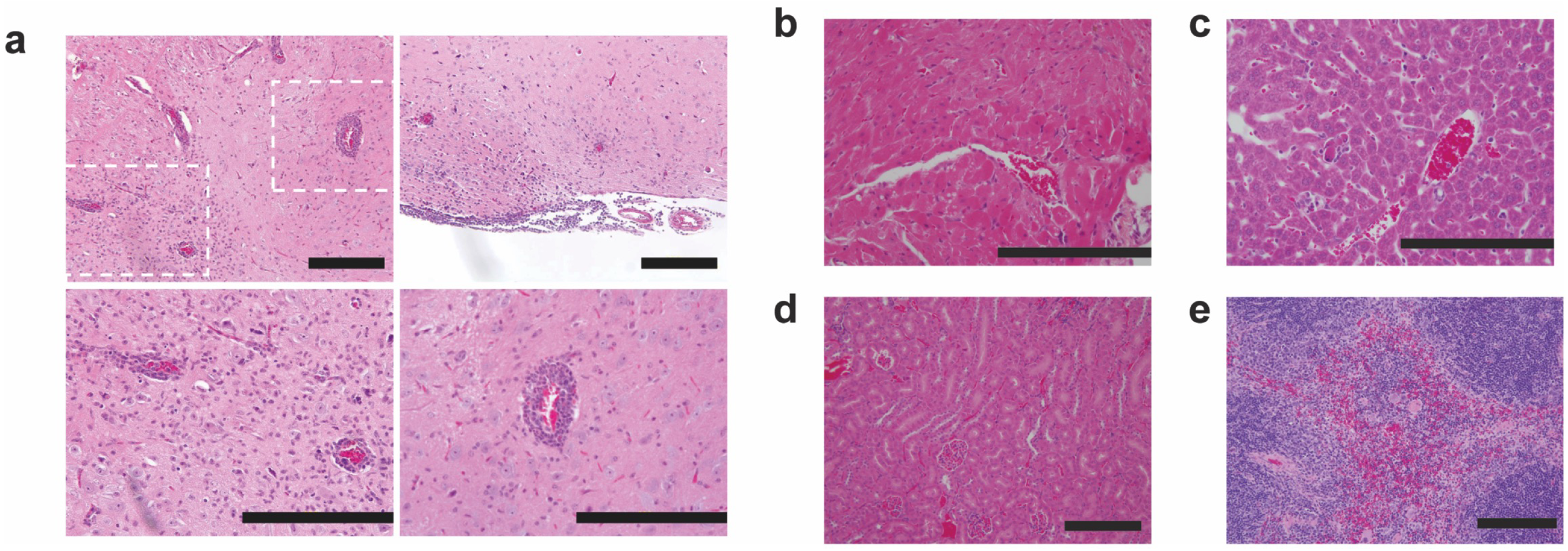
Extra-pulmonary histopathology after SARS-CoV-2 infection. **a-d**. Hematoxylin and eosin staining of the brain (**a**), heart (**b**), liver (**c**), kidney (**d**), and spleen (**e**) from K18-hACE2 mice following mock infection or at 7 dpi. Scale bars indicate 200 μm. For **a**, microscopic images show inflamed vessels with extravasation of immune cells into the brain parenchyma, microglial activation, and subarachnoid inflammation with involvement of the underlying parenchyma. The dashed box indicates the location of two higher power magnification images below.

**Extended Data Figure 4.**
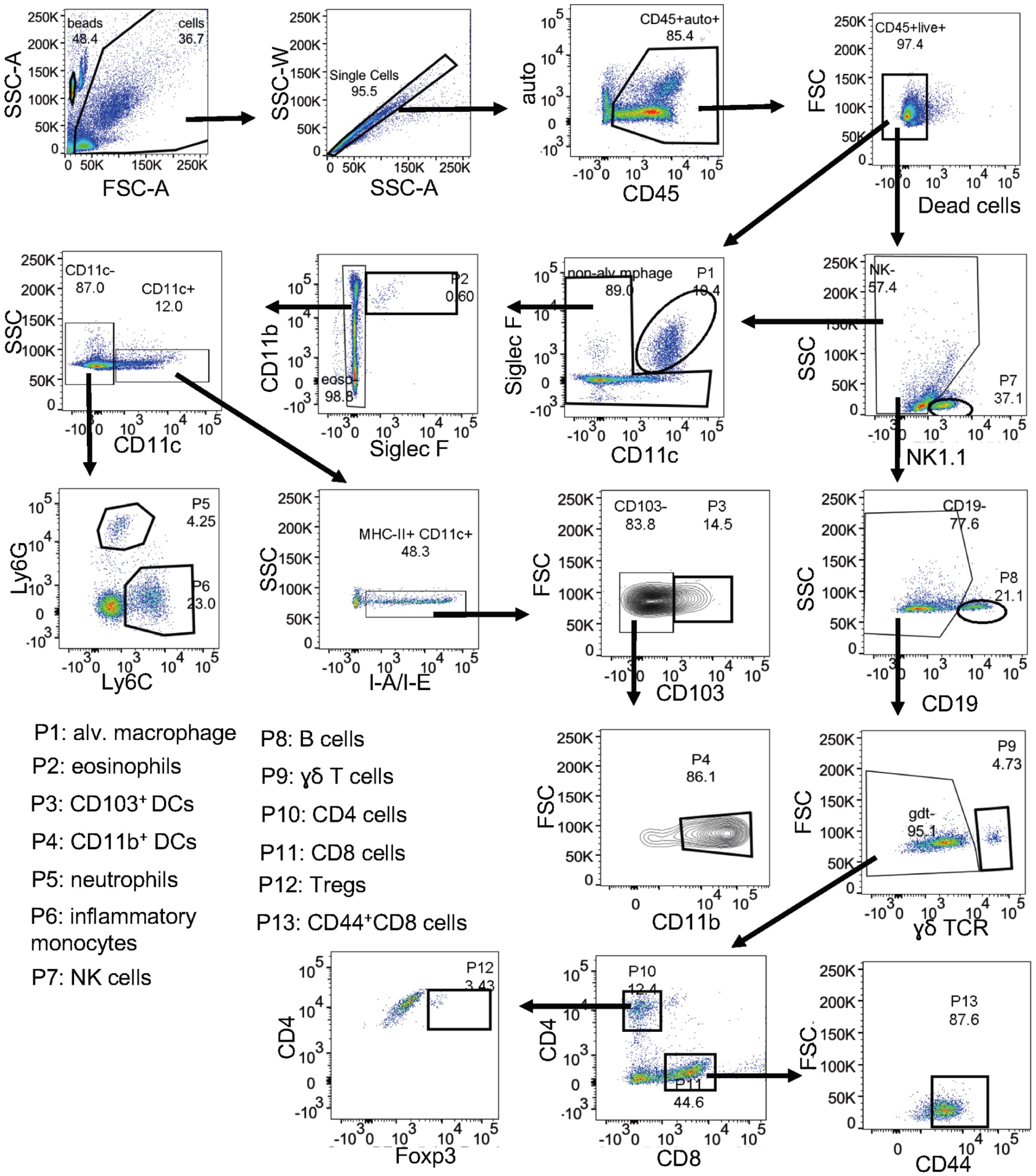
Flow cytometric analysis. Flow cytometric gating strategy for BAL and lung tissue analysis.

**Extended Data Figure 5.**
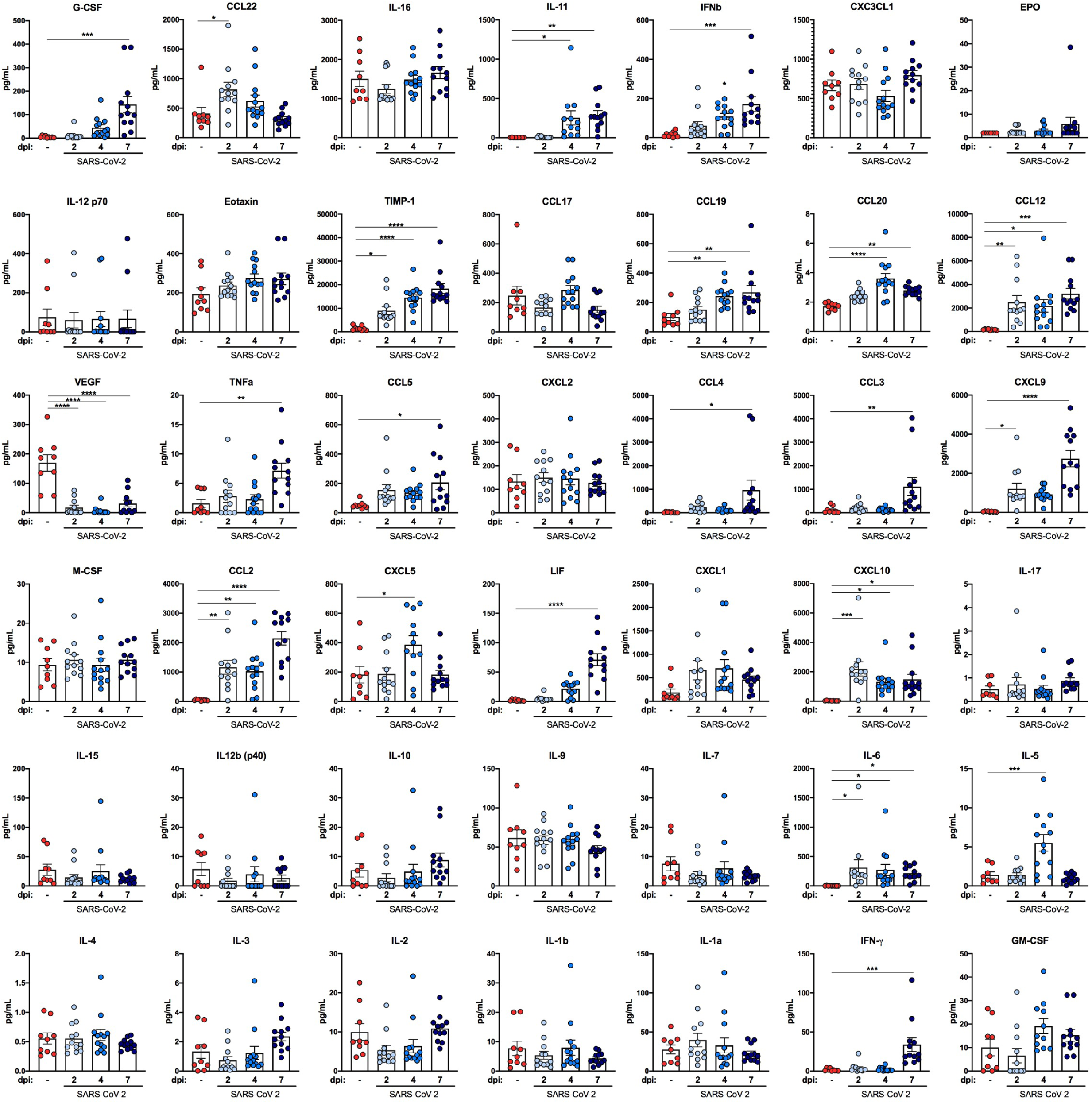
Cytokine induction following SARS-CoV-2 Infection. Cytokine levels as measured by multiplex platform in lung tissue of SARS-CoV-2-infected mice at 2, 4, and 7 dpi (two experiments, n = 9-11 per group; one-way ANOVA with Dunnett’s test; * *P* < 0.05; ** *P* < 0.01; *** *P* < 0.001, bars represent the mean value of each group).

**Extended Data Figure 6.**
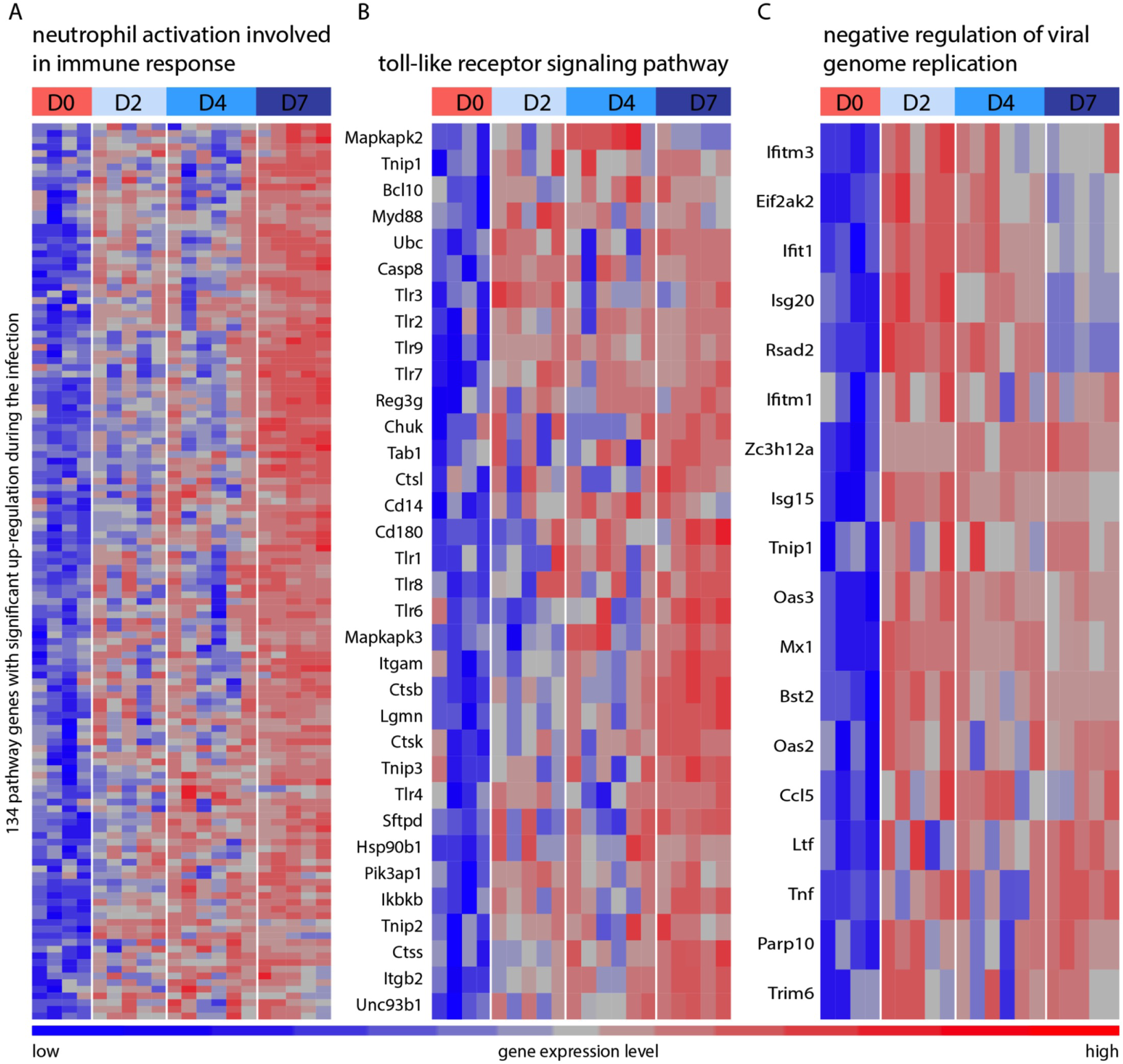
Transcriptional immune signatures following SARS-CoV-2 infection. **a-c**. Heat maps of significantly upregulated genes during SARS-CoV-2 infection enriched in neutrophil activation pathways (**a**), Toll-like receptor signaling pathway (**b**), and negative regulation of viral genome replication (**c**) identified through Gene Ontology analysis. Genes shown in each pathway are the union of differentially expressed genes from the three comparisons (2, 4, and 7 dpi versus mock-infected). Columns represent samples and rows represent genes. Gene expression levels in the heat maps are z score-normalized values determined from log2 [cpm values]).

**Supplementary Table 1**. Lists of up-regulated genes enriched in cytokine-mediated signaling pathway, type I IFN signaling pathway, cellular response to IFNγ, neutrophil activation pathways, toll-like receptor signaling pathways, and negative regulation of viral genome replication identified through Gene Ontology analysis and their associated q-value and fold-change values.

## METHODS

### Cells and viruses

Vero E6 (CRL-1586, American Type Culture Collection (ATCC), Vero CCL81 (ATCC), and Vero-furin cells^42^ were cultured at 37°C in Dulbecco’s Modified Eagle medium (DMEM) supplemented with 10% fetal bovine serum (FBS), 10 mM HEPES pH 7.3, 1 mM sodium pyruvate, 1× non-essential amino acids, and 100 U/ml of penicillin–streptomycin. The 2019n-CoV/USA_WA1/2019 isolate of SARS-CoV-2 was obtained from the US Centers for Disease Control (CDC). Infectious stocks were grown by inoculating Vero CCL81 cells and collecting supernatant upon observation of cytopathic effect; debris was removed by centrifugation and passage through a 0.22 μm filter. Supernatant was then aliquoted and stored at -80°C.

### Biosafety

All aspects of this study were approved by the office of Environmental Health and Safety at Washington University School of Medicine prior to the initiation of this study. Work with SARS-CoV-2 was performed in a BSL-3 laboratory by personnel equipped with powered air purifying respirators.

### Mice

Animal studies were carried out in accordance with the recommendations in the Guide for the Care and Use of Laboratory Animals of the National Institutes of Health. The protocols were approved by the Institutional Animal Care and Use Committee at the Washington University School of Medicine (assurance number A3381–01). Virus inoculations were performed under anesthesia that was induced and maintained with ketamine hydrochloride and xylazine, and all efforts were made to minimize animal suffering.

Heterozygous K18-hACE c57BL/6J mice (strain: 2B6.Cg-Tg(K18-ACE2)2Prlmn/J) were obtained from The Jackson Laboratory. Animals were housed in groups and fed standard chow diets. Mice of different ages and both sexes were administered 2.5 × 10^4^ PFU of SARS-CoV-2 via intranasal administration.

### Plaque forming assay

Vero-furin cells^42^ were seeded at a density of 2.5×10^5^ cells per well in flat-bottom 12-well tissue culture plates. The following day, media was removed and replaced with 200 μL of 10-fold serial dilutions of the material to be titered, diluted in DMEM+2% FBS. One hours later, 1 mL of methylcellulose overlay was added. Plates were incubated for 72 hours, then fixed with 4% paraformaldehyde (final concentration) in phosphate-buffered saline for 20 minutes. Plates were stained with 0.05% (w/v) crystal violet in 20% methanol and washed twice with distilled, deionized H20.

### Measurement of viral burden and *hACE2* expression

Tissues were weighed and homogenized with zirconia beads in a MagNA Lyser instrument (Roche Life Science) in 1000 μL of DMEM media supplemented with 2% heat-inactivated FBS. Tissue homogenates were clarified by centrifugation at 10,000 rpm for 5 min and stored at -80°C. RNA was extracted using the MagMax mirVana Total RNA isolation kit (Thermo Scientific) on the Kingfisher Flex extraction robot (Thermo Scientific). RNA was reverse transcribed and amplified using the TaqMan RNA-to-CT 1-Step Kit (ThermoFisher). Reverse transcription was carried out at 48°C for 15 min followed by 2 min at 95°C. Amplification was accomplished over 50 cycles as follows: 95°C for 15 s and 60°C for 1 min. Copies of SARS-CoV-2 N gene RNA in samples were determined using a previously published assay^8^. Briefly, a TaqMan assay was designed to target a highly conserved region of the N gene (Forward primer: ATGCTGCAATCGTGCTACAA; Reverse primer: GACTGCCGCCTCTGCTC; Probe: /56-FAM/TCAAGGAAC/ZEN/AACATTGCCAA/3IABkFQ/). This region was included in an RNA standard to allow for copy number determination down to 10 copies per reaction. The reaction mixture contained final concentrations of primers and probe of 500 and 100 nM, respectively.

For *hACE2* expression, RNA was DNase-treated (Thermo Scientific) following the manufacturer’s protocol. RNA levels were quantified as described above with the primer/probe set for *hACE2* (IDT assay: Hs.PT.58.27645939), compared to an RNA standard curve, and normalized to mg of tissue.

### Cytokine and chemokine mRNA measurements

RNA was isolated from lung homogenates as described above. cDNA was synthesized from DNAse-treated RNA using the High-Capacity cDNA Reverse Transcription kit (Thermo Scientific) with the addition of RNase inhibitor following the manufacturer’s protocol. Cytokine and chemokine expression was determined using TaqMan Fast Universal PCR master mix (Thermo Scientific) with commercial primers/probe sets specific for *IFN-g* (IDT: Mm.PT.58.41769240), *IL-6* (Mm.PT.58.10005566), *IL-1b* (Mm.PT.58.41616450), *TNF-a* (Mm.PT.58.12575861), *CXCL10* (Mm.PT.58.43575827), *CCL2* (Mm.PT.58.42151692), *CCL5* (Mm.PT.58.43548565), *CXCL11* (Mm.PT.58.10773148.g), *IFN-b* (Mm.PT.58.30132453.g), and *IL-28a/b* (Thermo Scientific Mm04204156_gH) and results were normalized to *GAPDH* (Mm.PT.39a.1) levels. Fold change was determined using the 2^-ΔΔCt^ method comparing treated mice to naïve controls.

### Cytokine and chemokine protein measurements

Lung homogenates were incubated with Triton-X-100 (1% final concentration) for 1 h at room temperature to inactivate SARS-CoV-2. Homogenates then were analyzed for cytokines and chemokines by Eve Technologies Corporation (Calgary, AB, Canada) using their Mouse Cytokine Array / Chemokine Array 44-Plex (MD44) platform.

### Histology and RNA *in situ* hybridization

Animals were euthanized before harvest and fixation of tissues. The left lung was first tied off at the left main bronchus and collected for viral RNA analysis. The right lung then was inflated with ∼1.2 mL of 10% neutral buffered formalin using a 3-mL syringe and catheter inserted into the trachea. For harvesting of brains for fixation, the mouse was decapitated, and the skull cap removed. The whole brain was removed intact, cut mid-sagittally to increase surface area of fixation, and drop fixed in 10% neutral-buffered formalin (NBF). For kidney, spleen, liver, and heart, organs were removed and drop-fixed in 10% NBF. For fixation after infection, organs were kept in a 40-mL suspension of NBF for 7 days before further processing. Tissues were embedded in paraffin, and sections were stained with hematoxylin and eosin. RNA *in situ* hybridization was performed using the RNAscope 2.5 HD Assay (Brown Kit) according to the manufacturer’s instructions (Advanced Cell Diagnostics). Briefly, sections were deparaffinized, treated with H_2_O_2_ and Protease Plus prior to probe hybridization. Probes specifically targeting hACE2 (cat no. 848151) or SARS-CoV-2 S sequence (cat no 848561) were hybridized followed by proprietary signal amplification and detection with 3,3’-Diaminobenzidine. Tissues were counterstained with Gill’s hematoxylin. An uninfected mouse was used as a negative control and stained in parallel. Tissue sections were visualized using a Nikon Eclipse microscope equipped with an Olympus DP71 camera, a Leica DM6B microscope equipped with a Leica DFC7000T camera, or an Olympus BX51 microscope with attached camera.

### Flow cytometry analysis of immune cell infiltrates

For analysis of BAL fluid, mice were sacrificed by ketamine overdose, followed by cannulation of the trachea with a 19-G canula. BAL was performed with three washes of 0.8 ml of sterile PBS. BAL fluid was centrifuged, and single cell suspensions were generated for staining. For analysis of lung tissue, mice were perfused with sterile PBS and the right inferior lung lobes were digested at 37°C with 630 µg/ml collagenase D (Roche) and 75 U/ml DNase I (Sigma) for 2 hours. Single cell suspensions of BAL and lung digests were preincubated with Fc Block antibody (BD PharMingen) in PBS + 2% heat-inactivated FBS for 10 min at room temperature before staining. Cells were incubated with antibodies against the following markers: AF700 anti-CD45 (clone 30 F-11), APC-Cy7 anti-CD11c (clone N418), PE anti-Siglec F (clone E50-2440; BD), PE-Cy7 anti-Ly6G (clone 1A8), BV605 anti-Ly6C (clone HK1.4; Biolegend), BV 711 anti-CD11b (clone M1/70), APC anti-CD103 (clone 2E7; eBioscience), PB anti-CD3 (clone 17A2), PE-Cy7, APC anti-CD4 (clone RM4-5), PE-Cy7 anti-CD8 (clone53-6.7), anti-NK1.1 (clone PK136), and BV605 anti-TCR γ/δ (clone GL3). All antibodies were used at a dilution of 1:200. Cells were stained for 20 min at 4°C, washed, fixed and permeabilized for intracellular staining with Foxp3/Transcription Factor Staining Buffer Set (eBioscience) according to manufacturer’s instructions. Cells were incubated overnight at 4°C with PE-Cy5 anti-Foxp3 (clone FJK-16s), washed, re-fixed with 4% PFA (EMS) for 20 min and resuspended in permeabilization buffer. Absolute cell counts were determined using TruCount beads (BD). Flow cytometry data were acquired on a cytometer (BD-X20; BD Biosciences) and analyzed using FlowJo software (Tree Star).

### Clinical laboratory analysis

Testing was performed on fresh whole-blood samples within a biosafety cabinet using point-of-care instruments. Prothrombin time was measured using the Coagucheck (Roche) meter. Electrolyte, acid-base, and hematology parameters were assayed on lithium-heparinized whole blood using the iSTAT-1 (Abbot) with the Chem8+ cartridge.

### Respiratory mechanics

Mice were anesthetized with ketamine/xylazine (100 mg/kg and 10 mg/kg, i.p., respectively). The trachea was isolated via dissection of the neck area and cannulated using an 18-gauge blunt metal cannula (typical resistance of 0.18 cmH_2_O.s/mL), which was secured in place with a nylon suture. The mouse then was connected to the flexiVent computer-controlled piston ventilator (SCIREQ Inc.) via the cannula, which was attached to the FX adaptor Y-tubing. Mechanical ventilation was initiated, and mice were given an additional 100 mg/kg of ketamine and 0.1 mg/mouse of the paralytic pancuronium bromide via intraperitoneal route to prevent breathing efforts against the ventilator and during measurements. Mice were ventilated using default settings for mice, which consisted in a positive end expiratory pressure at 3 cm H_2_O, a 10 mL/kg tidal volume (Vt), a respiratory rate at 150 breaths per minute (bpm), and an fraction of inspired oxygen (FiO_2_) of 0.21 (*i.e*., room air). Respiratory mechanics were assessed using the forced oscillation technique, as previously described^43^, using the latest version of the flexiVent operating software (flexiWare v8.1.3). Pressure-volume loops and measurements of inspiratory capacity were also done.

### Treadmill stress test

A six-lane mouse treadmill (Columbus Instruments, Columbus OH) was placed within a biosafety cabinet within the ABSL-3 laboratory. Mice were introduced to the treadmill test three times prior to infection, with each introductory session performed at increasingly faster rates. In general, the treadmill was set to ramp from 0 to maximum speed over the course of the first minute, then maintain maximum speed for 5 min. Failure to maintain adequate speed resulted in delivery of a shock; this occurred until the animal reinguaged the treadmill for a maximum of 5 failures. For each sex, we identified a speed at which >80% of mice successfully completed the test prior to infection (16 m/s for female; 14 m/s for male).

### RNA sequencing

cDNA libraries were constructed starting with 10 ng of total RNA from lung tissues of each sample that was extracted using a MagMax mirVana Total RNA isolation kit (Thermo Scientific). cDNA was generated using the Seqplex kit (Sigma-Aldrich) with amplification of 20 cycles. Library construction was performed using 100 ng of cDNA undergoing end repair, A tailing, ligation of universal TruSeq adapters and amplification of 8 cycles to incorporate unique dual index sequences. Libraries were sequenced on the NovaSeq 6000 (Illumina, San Diego, CA) targeting 40 million read pairs and extending 150 cycles with paired end reads. RNA-seq reads were aligned to the mouse Ensembl GRCh38.76 primary assembly and SARS-CoV-2 NCBI NC_045512 Wuhan-Hu-1 genome with STAR program (version 2.5.1a)^44^. Gene counts were derived from the number of uniquely aligned unambiguous reads by Subread:featureCount (version 1.4.6-p5)^45^. The ribosomal fraction, known junction saturation, and read distribution over known gene models were quantified with RSeQC (version 2.6.2)^46^. All gene counts were preprocessed with the R package EdgeR^47^ to adjust samples for differences in library size using the trimmed mean of M values (TMM) normalization procedure. Ribosomal genes and genes not expressed at a level greater than or equal to 1 count per million reads in the smallest group size were excluded from further analysis. The R package limma^48^ with voomWithQualityWeights function^49^ was utilized to calculate the weighted likelihoods for all samples, based on the observed mean-variance relationship of every gene and sample. Differentially expressed genes were defined as those with at least 2-fold difference between two individual groups at the Benjamini-Hochberg false-discovery rate (FDR) adjusted p-value, i.e. q-value < 0.05.

### Statistical analysis

Statistical significance was assigned when *P* values were < 0.05 using Prism Version 8 (GraphPad) and specific tests are indicated in the Figure legends. Analysis of weight change was determined by two-way ANOVA. Changes in functional parameters or immune parameters were compared to mock-infected animals and were analyzed by one-way ANOVA or one-way ANOVA with Dunnett’s test.

### Data availability

All data supporting the findings of this study are found within the paper and its Extended Data Figures, and are available from the corresponding author upon request. RNA sequencing data sets generated in this study are available at GEO: GSE154104.

